# In an “ecological-belonging” intervention to reduce inequities in STEM, context matters

**DOI:** 10.1101/2021.06.02.446772

**Authors:** Sarah P. Hammarlund, Cheryl Scott, Kevin R. Binning, Sehoya Cotner

## Abstract

Doubts about belonging in the classroom are often shouldered disproportionately by students from historically marginalized groups, which can lead to underperformance. *Ecological-belonging interventions* use a classroom-based activity to instill norms that adversity is normal, temporary and surmountable. Building on prior studies, we sought to identify the conditions under which such interventions are effective. In a chemistry course (Study 1), students from underrepresented ethnic backgrounds underperformed relative to their peers in the absence of the intervention. This performance gap was eliminated by the intervention. In an introductory biology course (Study 2), there were no large performance gaps in the absence of the intervention, and the intervention had no effect. Study 2 also explored the role of the instructor that delivers the intervention. The intervention boosted scores in classrooms of instructors with a fixed (versus growth-oriented) intelligence mindset. Our results suggest that ecological-belonging interventions are more effective in more threatening classroom contexts.

## Introduction

University-level science, technology, engineering and mathematics (STEM) courses often fail to provide an equitable experience for all students, leading to the exclusion of students from historically underserved groups (President’s Council of Advisors on Science and Technology 2012, National Research Council 2018). Students from underrepresented racial and ethnic backgrounds, women, and first-generation college students often perform lower than their peers in STEM courses, even when analyses correct for differences in pre-course preparedness (Lohfink & Paulsen 2005, Hyde et al. 2008, Matz et al. 2017, Salehi et al. 2019, Salehi et al. 2020). Among other factors, classroom social contexts—the daily interactions that students have with their peers, teaching assistants, and instructors—may contribute to disparities. For example, by virtue of their exposure to American society, most students and instructors in US classrooms are aware of the stereotypes that pertain to particular groups in academic settings (Steele 1997). When stereotypes are “in the air,” they may create psychological threat among negatively stereotyped students (e.g. stereotype threat; Steele & Aronson 1995, Steele 1997), which can impair their learning (Taylor & Walton 2011) and hinder their high-stakes test performance (Schamder & Sedikides 2018, Salehi et al. 2019). Importantly, non-stereotyped group members also commonly experience threat responses during interactions with negatively stereotyped others, particularly when they first meet (e.g., Blascovich et al. 2001, Goff et al. 2008). Perhaps unsurprisingly, students from negatively stereotyped groups tend to report lower levels of confidence and sense of belonging in science, compared to their non-stereotyped counterparts (Walton & Cohen 2007, Rainey et al. 2018, Gopalan & Brady 2020). Negative affective experiences can have an adverse impact on participation, performance, and persistence (Cohen & Garcia 2008, Schmader & Sedikides 2018). In particular, a low sense of belonging can affect how students interpret and respond to adversity (Wheeler & Petty 2001, Cohen & Garcia 2008). For example, if a student receives a low exam score, they may interpret the score as confirmation that “people like them” cannot succeed in the course and do not belong in STEM. This can lead to a self-fulfilling cycle of low performance and low sense of belonging that may result in withdrawal from STEM (Cohen & Garcia 2008, Rainey et al. 2019).

Recent work has attempted to negate the power of stereotypes in the classroom by changing classroom norms about the meaning and implications of adversity (e.g., when students struggle to learn difficult concepts or receive a poor exam grade). One recent study employed an *ecological-belonging intervention* that was designed to provide an alternative social norm within the classroom, instilling the idea that all students experience adversity and that challenges are normal, temporary and surmountable (Binning et al. 2020). The approach was developed from prior social-belonging interventions (Walton & Cohen, 2007; 2011, Walton et al. 2015, Yeager et al., 2016, Murphy et al. 2020) with the alteration that it is implemented in the classroom with discussion between instructors and peers in an effort to change the classroom “ecology” and intersubjective norms.

The roughly 30-minute intervention exercise consists of reflective writing about challenges that students anticipate encountering in the course, reading of testimonials from more advanced students (students from stereotyped and non-stereotyped backgrounds) who overcame past challenges, and a classroom discussion with peers and the instructor that was designed to reinforce and establish the intervention message as a local norm. Classroom discussions may be especially powerful for creating a common understanding that everyone struggles, regardless of their gender or ethnicity, which may rob factors like stereotype threat of their power. Binning et al. (2020) found that an ecological-belonging intervention improved performance of historically underperforming students. Specifically, they found improvement in course grades for ethnic minorities in an introductory biology course and for women in an introductory physics course.

Although these findings are promising, there are well-known concerns about the replicability of psychological science (Open Science Collaboration 2015) and social science more broadly (Camerer et al. 2018). Many high-profile studies have failed to replicate when studied in new contexts. Social-psychological interventions in education are no exception, as this work has yielded inconsistent replications (Walton 2014, Schwartz et al. 2016) and findings suggest that social context may powerfully moderate intervention effects (Walton 2014, Schwartz et al. 2016, Binning & Browman 2020, Walton & Yeager 2020).

The present research sought to identify the contexts in which an ecological-belonging intervention is most effective at promoting equity in college STEM classrooms. Across two studies, we trained current instructors and teaching assistants (TAs) to lead the intervention, most of whom taught two sections of the same course. Each instructor was then assigned to conduct the intervention in one of their class sections and to conduct their other section as usual with no changes. This design allowed for a strong test of the intervention’s robustness and potential for scalability beyond where it was first developed. It also allowed us to examine which features of the instructors’ course context are predictive of where the intervention may be effective. First, the results of Binning et al. (2020) suggest that the ecological-belonging intervention is most effective for students who experience underperformance—the intervention was effective where underperformance was occurring (e.g. women in physics) but not where it was not (e.g. women in biology). Second, a contextual factor that has yet to be explored is instructors’ beliefs about students’ intelligence. Instructors with so-called *growth mindsets* believe that intelligence is like a muscle that grows stronger with practice. Instructors with *fixed mindsets* believe that students are born with a certain amount of intelligence and cannot do anything to change their abilities (Dweck 2006, Dweck & Yeager 2019). Canning et al. (2019) recently showed that instructors’ mindsets about intelligence impacts students’ perceptions of themselves. In addition, Rainey et al. (2019) found that students who perceive that their instructors care about them feel a higher sense of belonging. Instructor mindset may therefore affect students’ sense of belonging, feelings of threat, and, in turn, the effectiveness of the intervention.

We report the results of ecological-belonging interventions conducted in two undergraduate STEM courses at a large public research university. Study 1 was conducted in an introductory chemistry lecture course for non-chemistry major students, and Study 2 in the laboratory sections associated with a non-majors introductory biology course. These courses had different structures and different performance gaps between students from various demographic groups. We sought to identify contexts in which the intervention is and is not effective at eliminating performance gaps between marginalized and non-marginalized students. Based on the results of Binning et al. (2020), we hypothesized that the intervention would only improve scores of students from demographic groups that showed performance gaps. We also hypothesized that the ecological-belonging intervention may be more effective for students of instructors with fixed mindsets, because those students may experience greater threat.

## Methods

### Study 1: Introductory Chemistry

#### Overview

Study 1 involved a non-majors lecture-based introductory chemistry course taught by one instructor with two sections. Each section had approximately 300 students. One section was randomly chosen to receive the intervention, and the other (“control” or “business-as-usual”) section was taught without any changes.

#### Intervention

The ecological-belonging intervention took place during two separate class periods spaced one week apart at the beginning of the Fall 2019 semester. During the first intervention activity, the instructor asked students to write a brief reflection about challenges that they anticipated facing during the course (see Supplementary Materials Section 1 for the intervention materials). The reflections were anonymous. These written responses were then collected. During the second intervention activity, the instructor presented quotes from students who had taken the course in the past. The quotes were adapted from Binning et al. 2020 and were intended to represent general concerns and challenges expressed by students. For example, a quote from Aniyah, a junior, read:

> “When I first got here, I was worried because it seemed like there weren’t many students like me. And I was really struggling with some of the chemistry concepts. It felt like everyone else was doing just fine, but I just wasn’t sure if I was cut out for the course. At some point during the first semester, I came to realize that, actually, a lot of other students were struggling, too. And I started to look at struggling as a positive thing. After I struggled with a hard problem and then I talked to other classmates and my TA about the solution—I realized that all that effort was worth it because it helped me learn and remember much more.”

Next, students were then asked to discuss these statements in small groups, using three prompts:

- *What are some common themes across several of the quotes we read?*
- *Why do you think that sometimes students don’t realize that other people are also struggling with the course?*
- *Why and how does people’s experience change over time? What do people do that helps them improve their experience with time?*

Finally, individual groups were called on to share some of their responses to the above prompts with the class, and the instructor facilitated a brief whole-class discussion.

#### Participants

A total of 610 students participated in Study 1. The control section had N = 271 students, and the intervention section had N = 339 students. However, due to missing data (see below) and students who withdrew at the start of the course, we analyzed data from N = 247 students in the control section and N = 303 students in the intervention section.

We examined four demographic variables: students’ gender, college generation, whether students belong to a minoritized and underrepresented ethinic or racial group, and whether students are of Asian descent. We use “male” and “female” to describe gender, but we recognize that these refer to biological sex rather than gender and may not represent how students identify. We used institutional data that unfortunately only include binary options. The total sample was 58% female and 42% male. Students were categorized as first-generation if neither of their parents attended college. The total sample was 18.5% first-generation and 81.5% continuing generation. Students from minoritized and under-represented ethnic or racial groups were categorized as “URM” (“under-represented minority”) (National Center for Science and Engineering Statistics 2019). URM status is based on institutional ethnicity data and includes students whose ethnicity is listed as American Indian, Black, Hawaiian, and Hispanic students. The total sample was 1.1% American Indian, 22% Asian, 5.5% Black, 0.33% Hawaiian, 2.8% Hispanic, 65% White, and 2.3% unknown. Asian and white students, who are over-represented in STEM relative to the general population, are designated as non-URM students. We note that “URM” is an imperfect designation. Some individuals in these groups may not identify with this term, and the term may hide differences in experiences of students of different ethnicities within this group. For the final demographic variable, we separated Asian students for analysis because while Asian students are represented in STEM at similar rates as in the general population (National Center for Science and Engineering Statistics 2019), Asian students may face unique challenges that white students do not face. All four demographic categories were equally represented in the control section compared to the intervention section (Gender: χ^2^ = 3.73, df = 1, p = 0.053; College generation: χ^2^ = 1.45, df = 1, p = 0.230; URM status: χ^2^ = 0.608, df = 1, p = 0.436; Asian status: χ^2^ = 2.88e-30, df = 1, p = 1).

#### Pre-course preparedness

We used grand mean-centered ACT and high school GPA scores as metrics of pre-course preparedness. There was no difference between students in the intervention and control sections in ACT and high school GPA scores (p >> 0.05). A minority of students were missing either ACT or high school GPA scores. We assigned those students the mean scores to avoid excluding them from the study.

#### Outcomes

The dependent variable we used was total course score, which is the number of course points obtained out of 100 possible percentage points at the end of the semester.

#### Data sources and missing data

We obtained demographic data from the university registrar’s student database, and course scores from the instructor. Individuals were anonymized by a researcher who was not associated with the study. In total, 75 students were excluded from analysis due to missing data. The majority of these students (67 students) were excluded because they dropped or withdrew from the course and therefore were missing total score data. The remaining students were excluded due to missing demographic data. Access to institutional data and grades were considered exempt from full review by the University’s Institutional Review Board (STUDY00000800). All students gave informed consent to participate in this research.

#### Statistical analysis

The analytic approach followed the approach employed by Binning et al. (2020). For both studies, analyses were performed using the Aiken and West (1991) procedure for probing for statistical interactions using multiple regression. Both studies used the same set of control variables (ACT and high school GPA scores). Study 1 analyses were performed using R version 3.6.0.

### Study 2: Introductory Biology

#### Overview

Study 2 involved a non-majors introductory biology course that largely serves students in the pre-health sciences and the natural sciences beyond biology. The course has both lecture and lab components. Three lecture sections run concurrently each semester, and each lecture section is associated with between 9 and 14 lab sections with maximum 24 students per section. The intervention took place in the lab sections during the first two weeks of the Fall 2019 semester. Lab sections (30 total) were led by graduate and undergraduate teaching assistants (TAs). A total of 16 TAs taught the lab sections. Of those, 12 TAs led two lab sections—for those TAs, one of their sections was randomly chosen to receive the intervention, and the other a control (“business-as-usual”) section. One TA led three lab sections, and two of them were randomly assigned to receive the intervention, and the third was assigned to be a control section. Two TAs taught only one section; both received the intervention. The remaining TA taught one intervention section and one section that was unknown—we disregarded that section. The intervention activities were identical to the procedure described above for Study 1.

#### Participants

A total of 588 students participated in Study 2. N = 324 students received the intervention, and N = 264 students were in control sections. As in Study 1, we used four demographic variables: gender (36% male, 64% female), college generation (76% continuing generation, 24% first-generation), URM status (80% non-URM, 16% URM, 4% unknown), and Asian status (80% non-Asian, 16% Asian, 4% unknown). Three demographic categories were equally represented in the control section compared to the intervention section (Gender: χ^2^ = 2.89, df = 1, p = 0.089; College generation: χ^2^ = 0.110, df = 1, p = 0.740; Asian status: χ^2^ = 0.068, df = 1, p = 0.790). URM students were overrepresented in the control sections (χ^2^ = 13.4, df = 1, p = 0.00026).

#### Teaching Assistant Mindset

The 16 TAs completed a survey including two Likert scale questions about their mindset about intelligence used by Canning et al. (2019). 1): Consider the undergraduate students you will teach and respond to this quote: “To be honest, students have a certain amount of intelligence, and they really can’t do much to change it,” and 2): Consider the undergraduate students you will teach and respond to this quote: “Your intelligence is something about you that you can’t change very much.” The responses to the two questions were averaged and mean-centered.

#### Pre-course preparedness

As in Study 1, we used students’ grand mean-centered ACT and high school GPA scores as a measure of pre-course preparedness. There was no difference between students in the intervention and control sections in ACT and high school GPA scores (p >> 0.05). A minority of students were missing either ACT or high school GPA scores. We assigned those students the mean scores in order to avoid excluding them from the study.

#### Outcomes

As in Study 1, the dependent variable we used was total course score, which is the number of course points obtained out of 100 possible percentage points.

#### Data sources and missing data

We obtained demographic data from the university registrar’s student database, and course scores from the instructors. Individuals were anonymized by a researcher who was not associated with the study. Analyses used restricted maximum likelihood estimation to handle missing data by using all available data (Raudenbush and Bryk 2002). However, runtime deletion excluded 12 students who were missing total course score data, bringing the analyzed model to N = 576. Access to institutional data and grades were considered exempt from full review by the University’s Institutional Review Board (STUDY00000800). All students and teaching assistants gave informed consent to participate in this research.

#### Statistical analysis

To account for the nestedness of the data, Study 2 employed a three-level hierarchical linear model (Raudenbush & Bryk 2002) using HLM 7.0 software. Students were divided into different lab sections, some sharing TAs, and some sharing lecture sections. The TAs (N = 16) were modeled at level 3, each TA’s sections (one treatment, one control; N = 32) were modeled at level 2, and students were modeled at level 1 (N = 588). The dependent variable, total course points, was in turn partially nested within one of three different lecture sections. As such, we entered two dummy codes at level 1 to account for differences in average performance across the three lecture sections. Also at level 1, we controlled for pre-course preparation by entering grand mean centered ACT scores and grand mean centered high school GPA. We also included dummy-coded variables to capture gender, first-generation, URM, and Asian demographic categories. At level 2, we included the classroom condition code (0 = control; 1 = treatment). In addition, to explore the potential contribution of TAs’ mindsets on their students’ performance, at level 3 we included TAs’ mean-centered growth mindset.

#### Data availability

Data and analysis scripts for both Study 1 and Study 2 are available in an online repository (Hammarlund et al. 2021).

## Results

### Study 1: Introductory Chemistry

#### Performance gaps

In order to understand whether certain students were underserved by the course, we examined scores in the control section to identify performance gaps. We identified demographic groups that underperformed relative to expectations based on pre-course preparedness metrics (high school GPA and ACT scores). Within the control section, under-represented ethnic and racial minority (“URM”) students scored 6.0 percentage points lower than non-URM (white and Asian) students (N = 247, B = -6.04, SE = 2.62, p = 0.022, t(234) = -2.30). We found no performance differences within other demographic categories (gender, college generation, and Asian/non-Asian students; Fig. 1).

**Figure 1.**
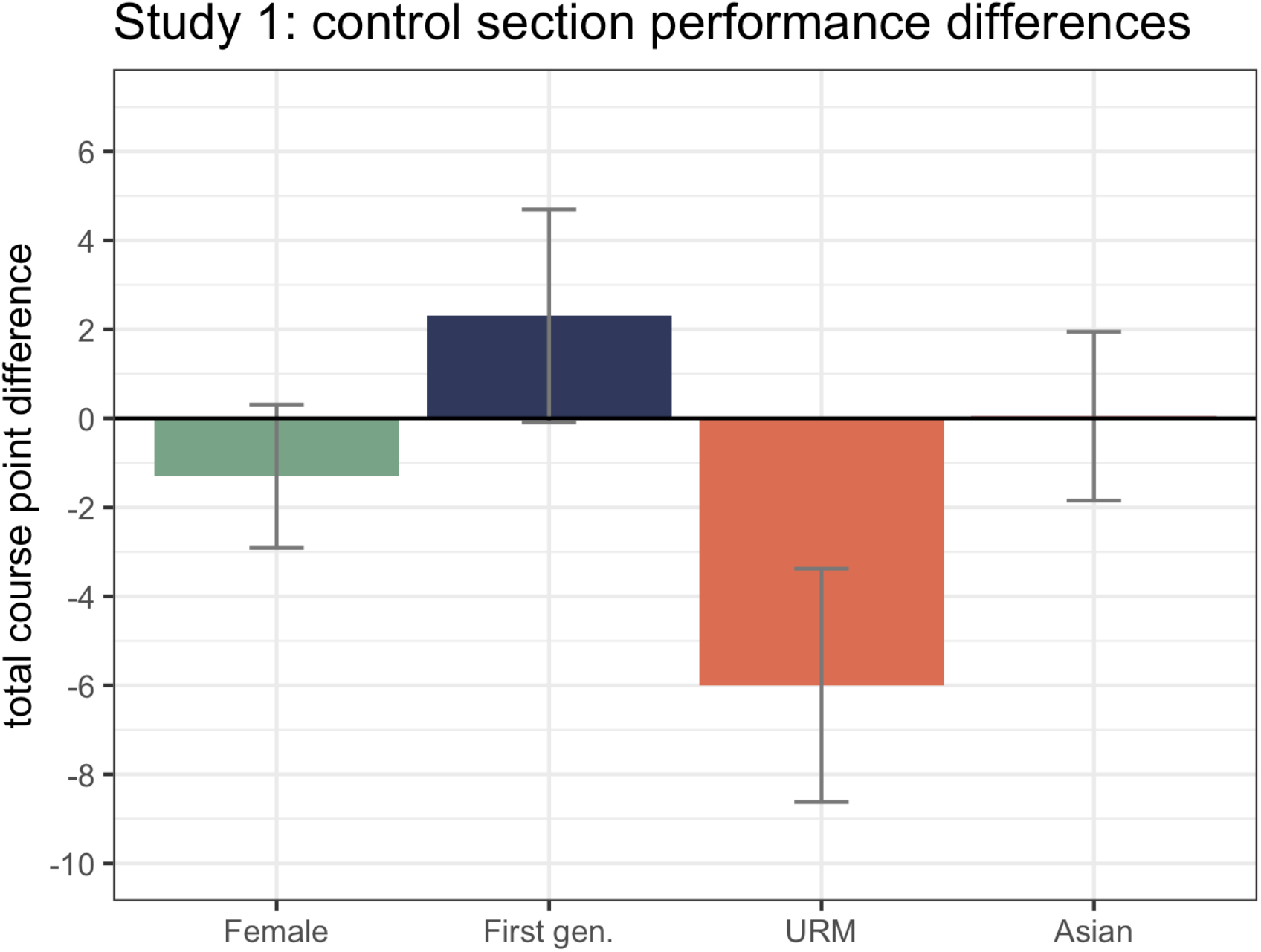
URM students are underserved in Introductory Chemistry. Analysis of total course points (out of 100) within the control (“business-as-usual”) section (N = 247). URM indicates under-represented racial or ethnic minority students, which encompasses American Indian, Hawaiian, Black and Hispanic students. The control section included 146 female students, 39 first-generation students, 28 URM students, and 57 Asian students. The y-axis value of 0 represents the performance of the reference group for each demographic group. For female students, the reference group is male students; for first-generation students, this is continuing generation students; for URM students, this is white and Asian students, and for Asian students, the reference group is students of all other ethnicities. URM students performed around 6 percentage points lower than white and Asian students (t(234) = -2.3, p = 0.022). Error bars show standard errors.

Because this analysis controlled for pre-course preparedness, the results indicate that underperformance emerged in the course among URM students, above and beyond prior differences in students’ pre-course preparedness. Although there are many reasons why performance gaps may emerge, we suggest that one factor may be that stereotypes were “in the air” (Steele 1997), which contributed to underperformance among URM students. We therefore hypothesized that the ecological-belonging intervention would reduce this performance gap seen in the control section.

#### Intervention outcomes

We found no main effect of the intervention on total course points (N = 550, B = 0.610, SE = 1.09, t(527) = 0.559, p = 0.576), indicating that the intervention had no effect on all students’ performance. However, we did find an interaction effect between URM status and the intervention, where URM students who received the intervention showed increased performance (Fig. 2; Supplementary Materials Section 2). Among non-URM (white and Asian) students, the intervention had no effect (B = -0.189, SE = 1.15, t(526) = -0.165, p = 0.869), but among URM students, the intervention was associated with an increase of 7.5 points (B = 7.54, SE = 3.36, t(526) = 2.24, p = 0.025), erasing the performance gap between white and Asian and URM students. There were no interaction effects with other demographic categories.

**Figure 2.**
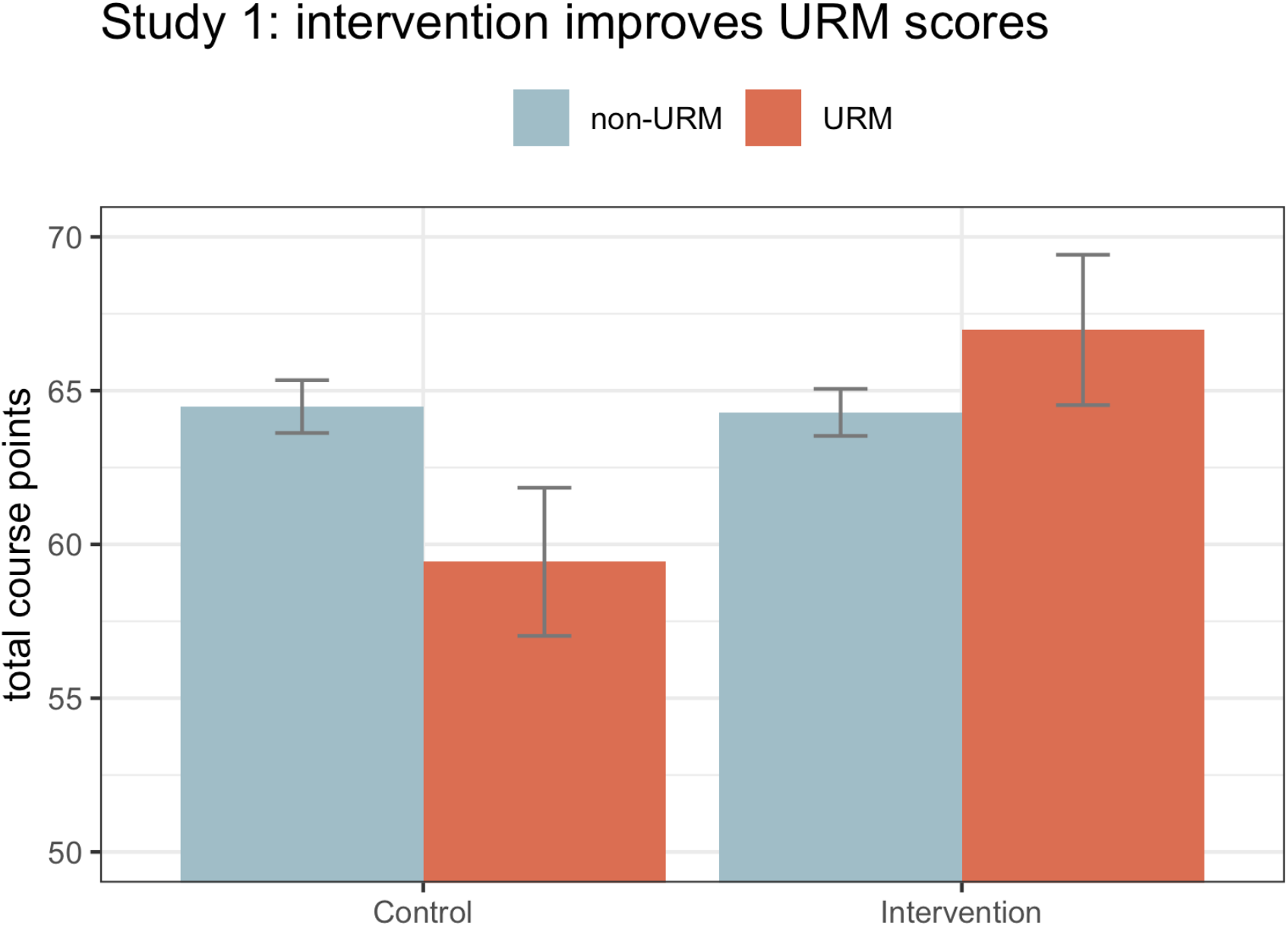
URM students benefit from the ecological-belonging intervention in Introductory Chemistry. Analysis shows an interaction effect between URM status and the intervention. Orange bars show URM students, and gray bars show non-URM (white and Asian) students. When URM students received the intervention, their scores increased by 7.54 points (t(526) = 2.242, p = 0.025), erasing the performance gap between URM and non-URM students. The control section contained 247 students total, with 28 URM students, and the intervention section contained 303 students total, with 27 URM students. Error bars show standard errors.

### Study 2: Introductory Biology

#### Performance gaps

As in Study 1, we first examined scores in the control sections to identify performance gaps. Within the control lab sections, first-generation college students performed 4.13 points lower than their continuing generation peers (t(251) = -2.36, p = 0.019; Fig. 3). We found no performance differences within other demographic categories (gender, URM, and Asian/non-Asian students). We hypothesized that first-generation students’ scores may improve as a result of the belonging intervention.

**Figure 3.**
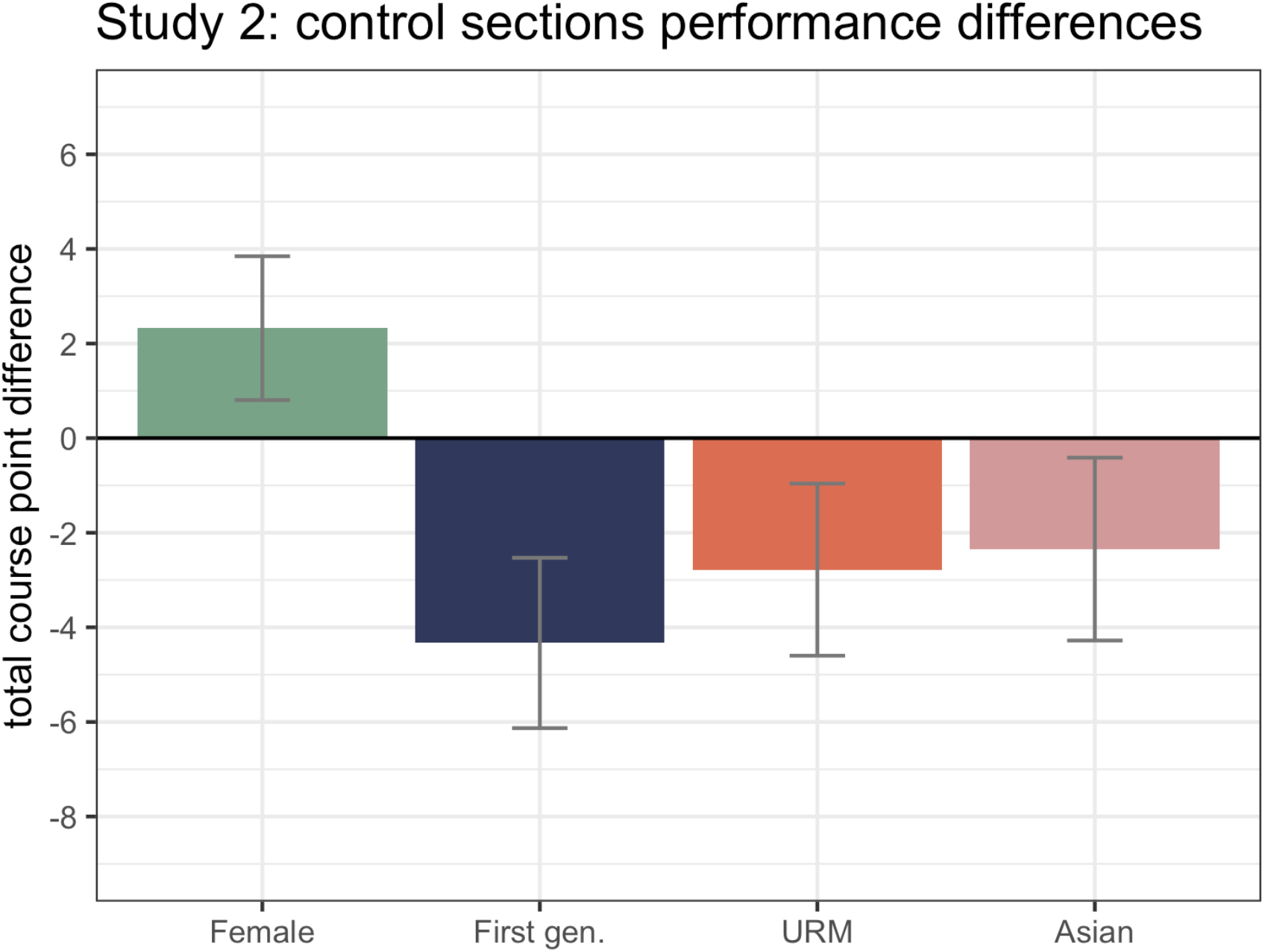
first-generation college students are underserved in Introductory Biology. Analysis of total course points (out of 100) of students within the control (“business-as-usual”) sections (N = 264). URM indicates under-represented racial or ethnic minority students, which encompasses American Indian, Hawaiian, Black and Hispanic students. The control sections included 179 female students, 66 first-generation students, 58 URM students, and 45 Asian students. The y-axis value of 0 represents the performance of the reference group for each demographic group. For female students, the reference group is male students; for first-generation students, this is continuing generation students; for URM students, this is white and Asian students, and for Asian students, the reference group is students of all other ethnicities. first-generation college students performed around 4 points lower than continuing generation students (t(251) = 1.75, p = 0.019). No other demographic categories showed significant performance differences. Error bars show standard errors.

**Figure 4.**
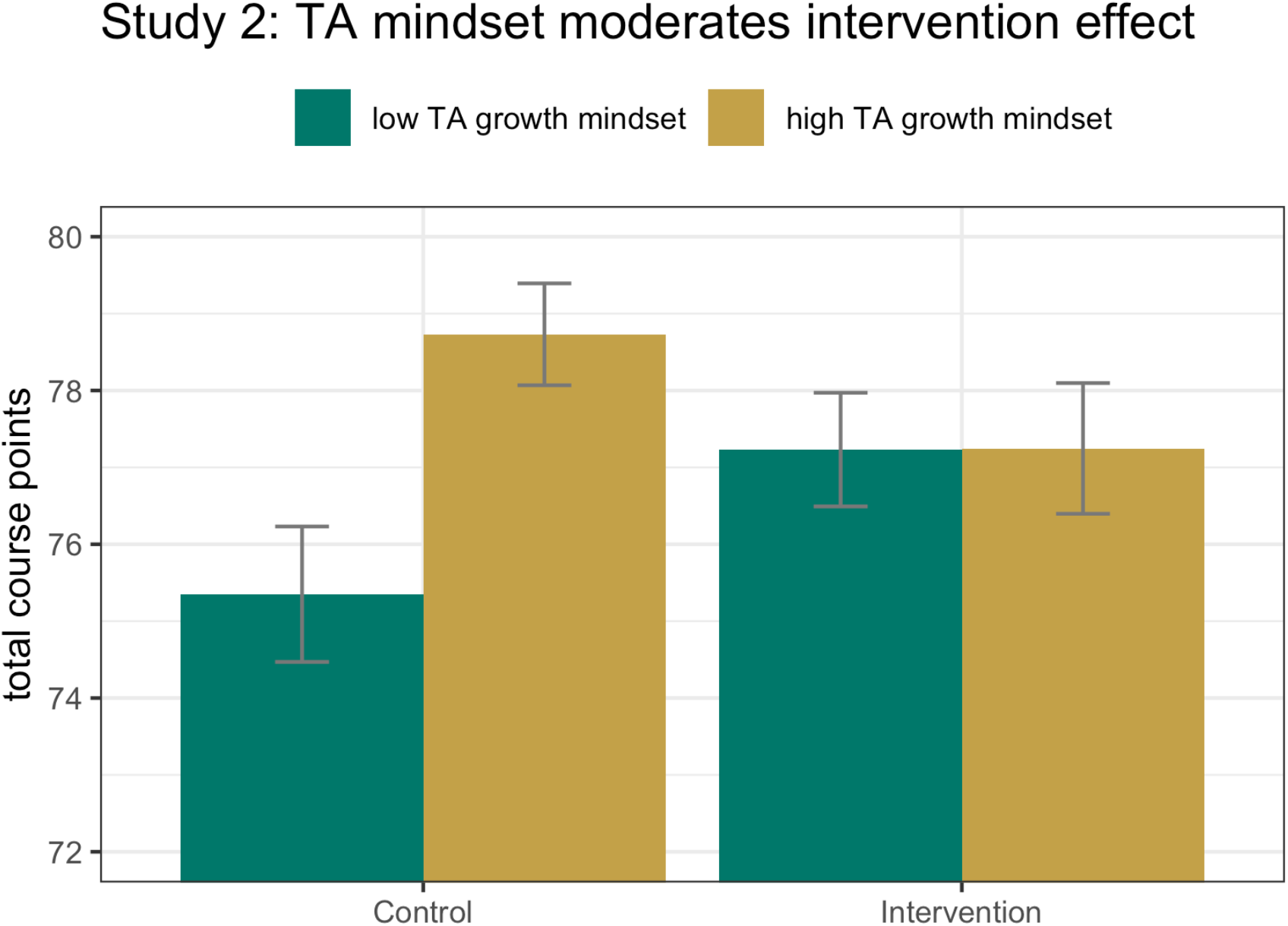
Teaching assistants’ views of students’ intelligence moderate the effects of the ecological-belonging intervention in Introductory Biology. TA mindset, a continuous variable, is shown as low (−1 standard deviation from the mean, green bars) and high (+1 standard deviation from the mean, gold bars). Within the control lab sections, students with TAs with a more growth-oriented mindset perform better than students of TAs with a more fixed mindset (t(14) = 3.02, p = 0.010). Within the intervention lab sections, TA mindset had no effect on student performance (t(14) = 0.06, p = 0.957). Students of TAs with more fixed mindsets (green bars) performed higher if they received the intervention (t(28) = 2.15, p = 0.04). Among students of TAs with more growth-oriented mindsets (gold bars), the intervention had no significant effect (t(28) = -1.34, p = 0.193). Error bars show standard errors.

#### Intervention outcomes

The test of the study hypotheses proceeded in several steps, with the first analyses attempting to directly replicate the effect seen in our Study 1 and in Study 1 of Binning et al. (2020), where the belonging intervention specifically improved scores of demographic groups that showed a performance gap. Analyses revealed that there was no overall main effect of the intervention on students’ performance at Level 2 (B = 0.20, SE = 0.93, t(28) = 0.22, p = 0.830; Supplementary Materials Section 3), nor was there a differential cross-level interaction between the intervention and first-generation status (B = -1.05, SE = 2.14, t(564) -0.49, p = 0.623). This means that first-generation college students’ scores did not improve as a result of the intervention. Furthermore, there was no cross-level interaction between the intervention and URM status (B = -2.28, SE = - 2.57, t(564) = -0.89, p = 0.376).

To explore the effect of TA growth mindset on students’ performance, and to examine if the intervention had effects among TAs with a low (versus high) growth mindset, we tested for a TA mindset (Level 3) x intervention (Level 2) interaction on students’ course grades. The analysis yielded a significant interaction (B = -1.44, SE = 0.36, t(28) = 4.01, p = 0.001). Analysis of the effect of the intervention across levels of TA mindset revealed that the intervention had a positive effect among TAs with a relatively low growth mindset (−1 SD; B = 2.01, SE = 0.93, t(28) = 2.15, p = 0.04). By contrast, the intervention had no effect in lab sections led by TAs with a strong growth mindset (+1 SD; B = -1.34, SE = 1.00, t(28) = -1.34, p = 0.193).

Notably, in the control classrooms, TA mindset was significantly associated with course scores (B = 1.47, SE = 0.49, t(14) = 3.02, p = 0.010). That is, TAs’ mindsets were predictive of their students’ course grades such that TAs with a strong growth-oriented mindset had students who performed better in the class compared to TAs with a relatively fixed mindset about intelligence. However, within classrooms where TAs delivered the belonging intervention, TA mindset had no effect on students’ scores (B = 0.02, SE = 0.41, t(14) = 0.06, p = 0.957).

## Discussion

We predicted that the intervention would help students in groups that underperformed, relative to their peers, in the control sections of the two courses. In other words, the intervention would only be effective for students who experience a sense of threat and uncertainty about belonging that leads to a performance gap. This is consistent with findings from similar interventions (Schwartz et al. 2016, Binning et al. 2020). In Study 1, where underrepresented ethnic and racial minority (URM) students underperformed relative to white and Asian students, we found that the intervention did improve URM students’ scores. In Study 2, URM students did not significantly underperform in the control condition. However, first-generation college students underperformed relative to continuing generation students. We found that the intervention had no effect on either URM or first-generation students.

In Study 2, we also predicted that instructors’ mindsets about intelligence would affect the effectiveness of the intervention. Having an instructor with a fixed mindset may increase the threat that students feel in the classroom, creating greater potential for the intervention to be effective. Indeed, in the control sections, students of instructors with more fixed mindsets had lower scores than students of instructors with more growth-oriented mindsets. We found that the intervention specifically improved scores of students whose teaching assistants had a fixed view of intelligence, but it had no effect on students whose teaching assistants had a growth mindset. When a TA with a fixed mindset performed the intervention, this may have changed students’ perceptions of their TA, making the TA’s theories of intelligence irrelevant. In addition, the intervention may have provided students with tools to counter negative signals from fixed mindset TAs. Students of TAs with high growth mindsets may have already felt less threat, and therefore did not stand to benefit from the intervention.

Overall, our results support the idea that ecological-belonging interventions are most effective in threatening contexts. The intervention bolstered scores of URM students in contexts where they were underperforming, but it had no effect in contexts where they were not. The intervention improved performance of students whose TA had a fixed mindset, but it had no effect when TAs had a growth mindset. In other words, the intervention appears to be effective only where it is needed. We also found no negative effects of the intervention, so while the intervention does not improve performance across the board, it does not *harm* students’ performance.

The results have promising practical implications. Importantly, this research demonstrates that the ecological-belonging intervention can be effectively exported to new university contexts, effectively taught to instructors, and effectively delivered with impact. Unlike Binning et al., (2020), the instructors who delivered the intervention were not discipline-based education researchers. In fact, many were undergraduate and graduate-level teaching assistants with limited experience in the classroom and little—if any—pedagogical training. This intervention training was brief and would be easy to implement at a larger scale (adaptable materials are provided in the Supplementary Materials Section 1). Indeed, we know of several instructors at other institutions around the US that plan to implement similar interventions, thus enabling us to further contextualize the conditions in which a belonging exercise is helpful. In addition, this is the first time ecological-belonging interventions have been implemented in non-majors courses and in courses taken mainly by second-through fourth-year students. This suggests that these interventions have power beyond “gateway” majors courses taken at the start of students’ university experiences.

An important caveat is that while a threatening classroom context may be necessary for the intervention to have an effect, it is not sufficient. The lack of intervention benefits for first-generation students in Study 2 illustrates this point. There was a small but significant performance gap between first-generation and continuing generation students in the control sections, but this gap was also present in the intervention sections. Previous research has shown that social-psychological interventions are capable of improving outcomes among first-generation college students (Harackiewicz et al. 2014, Stephens et al. 2014). Although it is difficult to speculate about the causes of null effects, we believe that one important distinction may be the relative visibility of students’ URM status versus their first-generation status. In particular, whereas URM status is commonly (though not always) visible (e.g., due to physical characteristics), first-generation status is usually not visible. Thus, when students were engaging in the intervention, they would not know whether a peer was a first-generation student unless that peer volunteered the information. If the intervention works by shaping intersubjective norms, perhaps a lack of visibility makes the intervention less effective for addressing less visible sources of belonging uncertainty. This is an important question for future research.

Several standard limitations of human-subjects research apply to our study. Specifically, participating students and TAs represented a range of backgrounds, experiences, and axes of diversity. We were unable to control for all sources of variety, some of which may have affected our findings. We were also unable to explore students’ intersecting identities, due to limited sample sizes. Additionally, we only investigated one response variable (overall performance), so we do not know what, if any, additional impacts the intervention may have had. Critically, we did not measure actual “sense of belonging” in either population. Furthermore, Study 1 had one intervention and one control section, limiting our ability to extrapolate beyond the study population. However, our findings resonate with those of prior work on belonging interventions (Binning et al. 2020) and instructor mindset (Canning et al. 2019), while also adding nuance to our current understanding of the impact of course-level interventions.

Our findings suggest several important avenues for future research. We began with reference to the concerns about the replicability of psychological science, and our findings confirm that results are dependent on the context where it is delivered (Van Bavel et al. 2016). Thus, we reiterate prior calls for further replication in other contexts (i.e. beyond large research institutions, in other disciplines, with different documented challenges). Future work could illustrate whether the intervention has a lasting effect. Related interventions have shown effects that persist over multiple years and varied social settings (e.g., after students graduate; Brady et al. 2020, Goyer et al. 2017), but the duration and generality of ecological-belonging interventions need further study.

In addition to refining our contextual understanding, future work should target a mechanistic understanding—*why* do these brief interventions work? Establishing mechanism will likely require a rigorous qualitative investigation; this could involve interviewing students at different timepoints to investigate how the intervention affected their sense of belonging (à la Rainey et al. 2018), confidence about overcoming adversity, awareness of other students’ struggles, feelings about whether other students like them and care about them, and feelings about whether the instructor cared about them. Using instruments that measure belonging (such as the Psychological Sense of School Membership Scale; Goodenow 1993) could also elucidate whether the mechanism by which we’re *assuming* the intervention works is accurate (Schwartz et al. 2016).

We also suggest that instructor characteristics beyond mindset may impact the intervention’s effectiveness. As more interventions are conducted and assessed, it will become possible to conduct a meta-analysis to explore the predictive power of other instructor characteristics on student outcomes. For example, does it matter if the instructor is the same race, ethnicity, or gender as the students experiencing threat? Similarly, can an intervention compensate for having an instructor that does not match a student’s identity?

Finally, while overseeing the interventions, we noted, anecdotally, that merely reading the concerns of their students (which we did not require) could impact TAs, leading them to express surprise at how worried the students were about the material, interacting with the instructor, or managing the course requirements. We are currently exploring the impact of the intervention on the TAs themselves, and we are motivated by one primary question: can delivering an intervention foster greater instructor empathy for students?

Our findings add to the current dialogue on both the promise and caveats of psycho-social interventions, specifically those designed to mitigate barriers to equity in STEM education. These interventions are not a panacea for all challenges to equity in STEM, but that does not invalidate their implementation. Rather, this work calls for continued documentation of the criteria under which belonging interventions are effective. Critically, any intervention should be envisioned as a short-term solution. Our collective goal as STEM educators should be to realize a future in which our scientists represent our current population, and belonging interventions are no longer necessary.

## Acknowledgements

This project was supported by the SEISMIC collaboration (www.seismicproject.org). Funding for the SEISMIC project has been provided by the Alfred P. Sloan Foundation and the Participating Institutions. This project was also supported by the Biology Teaching Assistant Project (BioTAP), an NSF-Funded Research Coordination Network (DBI 1539903, Principal investigator: Dr. Elisabeth Schussler). S.H was supported by a NSF Graduate Research Fellowship. We thank members of the Cotner lab for feedback and support, especially Dr. Sadie Hebert for assistance with data collection and curation.

## Author Biographical

Sarah Hammarlund is affiliated with the Department of Ecology, Evolution and Behavior and the BioTechnology Institute at the University of Minnesota. Cheryl Scott is affiliated with the Department of Biology Teaching and Learning at the University of Minnesota. Kevin Binning is affiliated with the Department of Psychology and the Learning Research and Development Center at the University of Pittsburgh. Sehoya Cotner is affiliated with the Department of Biology Teaching and Learning at the University of Minnesota and the Department of Biological Sciences at the University of Bergen.

