## Supplementary Materials for "In an “ecological-belonging” intervention to reduce inequities in STEM, context matters"

**Supplementary Materials for Hammarlund et al.**
***In an "ecological-belonging" intervention to reduce inequities in STEM, context matters***

#### Contents


**Section 1: Intervention Materials**

**1.1**: Lesson plan
**1.2**: Week 1 writing prompt
**1.3**: Student quotes

**Section 2: Study 1 Results**

**2.1**: Performance gaps analysis
**2.2**: Main effect of the intervention
**2.2**: Interaction effect between the intervention and URM status

**Section 3: Study 2 Results**

**3.1**: Performance gaps analysis
**3.2**: Main effect of the intervention and TA mindset
**3.3**: Interaction effect between the intervention and first generation status
**3.4**: Interaction effect between the intervention and TA mindset

### Section 1: Intervention Materials

#### 1.1. Lesson Plan

The following is the lesson plan given to teaching assistants (TAs) in Study 2. The instructor for the Study 1 course was given nearly identical materials.

**Belonging Mindset Classroom Discussion Activity for Biology 1009**

**ACTIVITY SCRIPT FOR TEACHING ASSISTANTS**

##### Week One

**Introduction 2 minutes**

TA introduces the writing activity: *"Before we get started, we are going to take a break from our regular lab activities to talk about Biology concerns. We are always trying to make things better for our students, and so today the Department of Biology Teaching and Learning is asking for your help to tell us a little about your concerns. For many of you, this is your first college-level biology course, it may be your first time away from home, you are meeting a lot of new people, taking on a lot of new courses, and trying to find your place here at the University of Minnesota. It can be easy to feel overwhelmed and to ask yourself, "Do I really belong here?" and "Am I smart enough to make it?"*

*These kinds of experiences are normal. Everyone goes through them, and they get better with time as you adjust.*

**Writing 8-10 minutes**

TA distributes paper with the writing prompt: *"Today, we'd like each of you to reflect on some of the concerns you may have about taking Biology 1009. What do you think will be difficult or challenging for you? These concerns may be about course content, navigating resources, working in groups, interacting with your TA or professor, and so on. Please take a few minutes to explain your thoughts on the sheet of paper I provided.*

*Please don't include your name or other identifying information in what you write. But please do write legibly.*

*When the ten minutes are up, I will collect what you've written. Please write on your own, individually. We'll return to this discussion next week."*

Notes for TA:

**What if…

**A student isn't writing?**

Try first asking (or nonverbally signing) to see if they are okay and understand the directions. If they are okay, do understand, and aren't being disruptive, let them work at their own pace.

**A student or group of students is being disruptive?**

First, try nonverbally signing to get them back on track. If that doesn't work, "Hey, we want everyone to be able to think and write, so it'd be helpful if you were [describe appropriate behavior].

At the end of 8-10 minutes, students' writing samples are collected.

#### Week Two

**Reading 5 minutes**

*"As I looked through a few examples of what you all have written, I see a lot of very common concerns that you have. I am also not surprised that I had some of the same concerns when I took general biology. In preparation for today's exercise, a team interviewed a number of juniors and seniors who described their experiences in Biology 1009. I'd like to take a few minutes to provide some examples that these students shared."*

TA then hands out the quotes for students to read.

**Small group discussion 10 minutes**

**4 students in each group.**

*"For the next ten minutes, I ask that you please discuss with the small group of 3-4 students sitting near you what you wrote about and the quotes you have just read. Please answer the following questions as a group:"* (TA writes these questions on the board/.ppt, may need to help students get into small groups).

- What are some common themes across several of the quotes we read?
- Why do you think that sometimes students don't realize that other people are also struggling with the course?
- Why and how does people's experience change over time? What do people do that helps them improve their experience with time?

"After a few minutes, I will call on each group to share some of their responses to these questions."

Notes for TA:

**What if…

**A student says something that is off-topic or inappropriate?**

Blink and continue. Don't contradict, but breeze past it. Consider reframing or drawing on personal experience.

**People aren't talking or we are starting to run out of things to say.**

Approach group and ask students "why" and "how":

- Why do you think people wonder at first if they belong?
- Why do you think people often think they're the only one who worries about whether they fit in in college?
- Ask students to share additional experiences that resonate with what the group has been discussing.

TA: Share your own experience that resonates with what the group has been discussing (if applicable).

**Closing 5-7 minutes**

Now the TA will ask the students to share their group discussion with the class.

*"All right, let's come back together. I've been overhearing some great discussions and I'd love to hear your thoughts."*

Solicit input on the three questions on the board, ultimately getting input from each group, even if it's just "does your group have anything to add?" or "did any of these student concerns seem similar to those you expressed last week?"

*Conclude with: "I think your feedback is going to be very helpful for the department. But my real hope is that these discussions may help you succeed in Biology 1009."*

#### 1.2. Writing Prompt

The following page contains the handout that students in Study 2 were given in week 1 of the intervention. The prompt for Study 1 was nearly identical.

Take some time to reflect on some of the concerns you may have about taking Biology 1009. What do you think will be difficult or challenging for you? These concerns may be about course content, navigating resources, working in groups, interacting with your TA or professor, and so on.

Don't include your name or other identifying information in what you write. But please do write legibly. We'll return to this discussion next week.

##### 1.3. Student Quotes

The following pages contain the quotes that students were given in week 2 of the intervention for Study 1 (chemistry), followed by Study 2 (biology). The acronyms after the students' names show the college they belong to. CBS = College of Biological Sciences, CFANS = College of Food, Agricultural and Natural Resource Sciences, CLA = College of Liberal Arts, CEHD = College of Education and Human Development, CSE = College of Science and Engineering.

**Quotes from former Chemistry 1061 students**

"I remember taking my first chemistry class as a freshman. Before coming to college, I didn't worry much about grades, so I felt unprepared for the increased workload and differences in grading. I remember being surprised after getting burned grade-wise several times, and feeling stressed as a result. But then I got some help from the instructor and the TA, found a study group, and was able to turn things around. Looking back now, I think my struggles were pretty normal. Even though people don't like to admit it, basically everyone has trouble with certain concepts. Although it was a somewhat rocky start, it felt good to learn from my mistakes, and I am proud of the success I have had."

-James, UMN Senior, CBS

When I first got to the university, it was overwhelming. I didn't know all the rules of navigating different resources, and sometimes I felt embarrassed to ask questions—so I didn't ask. However, I quickly learned that other students usually had the same question I did, and we all benefitted from working with each other and learning from each other. I've also gotten much better at advocating for myself. Sometimes I have difficulty with an idea that my classmates understand. Other times, they struggle with concepts that I understand. I remember there wasn't always an "aha!" moment, where everything clicked. It was usually much more gradual, with some concepts only becoming clear after lots of practice and discussion with my study group. I realized that everyone struggles some times, and the important thing is to not give up and help each other out."

-Julia, UMN Senior, CSE

"When I first got here, I was worried because it seemed like there weren't many students like me. And I was really struggling with some of the chemistry concepts. It felt like everyone else was doing just fine, but I just wasn't sure if I was cut out for the course. At some point during the first semester, I came to realize that, actually, a lot of other students were struggling, too. And I started to look at struggling as a positive thing. After I struggled with a hard problem and then I talked to other classmates and my TA about the solution—I realized that all that effort was worth it because it helped me learn and remember much more."

-Aniyah, Junior, CLA

"I was worried that my high school courses had not prepared me well for college. Honestly, when I got here, I thought professors were scary. I thought they were critical and hard in their grading, and sometimes it felt like they put things on the quizzes or exams that we hadn't discussed in class. But then I realized that the professor wanted me to be able to apply the concepts in many different situations. So I started to study in a way that would help me do that, and I did my best to learn from my mistakes on quizzes and exams. And I saw that even when the professors' grading seemed tough, it didn't mean they looked down on me or that I didn't belong. It was just their way of motivating high achieving students."

- Anil, Senior, CFANS

**Quotes from former Biology 1009 students**

"I remember taking my first biology class as a freshman. Before coming to college, I didn't worry much about grades, so I felt unprepared for the increased workload and differences in grading. I remember being surprised after getting burned grade-wise several times, and feeling stressed as a result. But then I got some help from the instructor and the TA, found a study group, and was able to turn things around. Looking back now, I think my struggles were pretty normal. Even though people don't like to admit it, basically everyone has trouble with certain concepts. Although it was a somewhat rocky start, it felt good to learn from my mistakes, and I am proud of the success I have had."

-James, UMN Senior, CBS

When I first got to the university, it was overwhelming. I didn't know all the rules of navigating different resources, and sometimes I felt embarrassed to ask questions—so I didn't ask. However, I quickly learned that other students usually had the same question I did, and we all benefitted from working with each other and learning from each other. I've also gotten much better at advocating for myself. Sometimes I have difficulty with an idea that my classmates understand. Other times, they struggle with concepts that I understand. I remember there wasn't always an "aha!" moment, where everything clicked. It was usually much more gradual, with some concepts only becoming clear after lots of practice and discussion with my study group. I realized that everyone struggles some times, and the important thing is to not give up and help each other out."

-Julia, UMN Senior, CFANS

"When I first got here, I was worried because it seemed like there weren't many students like me. And I was really struggling with some of the biology concepts. It felt like everyone else was doing just fine, but I just wasn't sure if I was cut out for the course. At some point during the first semester, I came to realize that, actually, a lot of other students were struggling, too. And I started to look at struggling as a positive thing. After I struggled with a hard problem and then I talked to other classmates and my TA about the solution—I realized that all that effort was worth it because it helped me learn and remember much more."

-Aniyah, Junior, CLA

"I was worried that my high school courses had not prepared me well for college. Honestly, when I got here, I thought professors were scary. I thought they were critical and hard in their grading, and sometimes it felt like they put things on the quizzes or exams that we hadn't discussed in class. But then I realized that the professor wanted me to be able to apply the biology concepts in many different situations. So I started to study in a way that would help me do that, and I did my best to learn from my mistakes on quizzes and exams. And I saw that even when the professors' grading seemed tough, it didn't mean they looked down on me or that I didn't belong. It was just their way of motivating high achieving students."

- Anil, Senior, CEHD

#### Section 2: Study 1 Results

##### 2.1. Performance gaps analysis

Note: control section only

| | Coefficient | Standard error | T-ratio | d.f. | P-value |
|---|---|---|---|---|---|
| Intercept | 64.1 | 0.793 | 80.8 | 234 | < 0.001 |
| Female | -1.30 | 1.61 | -0.807 | 234 | 0.420 |
| First-generation | 2.28 | 2.39 | 0.955 | 234 | 0.341 |
| URM | -6.04 | 2.62 | -2.30 | 234 | 0.022 |
| Asian | 0.051 | 1.90 | 0.027 | 234 | 0.979 |
| ACT score | 1.44 | 0.272 | 5.31 | 234 | < 0.001 |
| High school GPA | 8.42 | 2.73 | 3.09 | 234 | < 0.001 |

##### 2.2. Main effect of the intervention

| | Coefficient | Standard error | T-ratio | d.f. | P-value |
|---|---|---|---|---|---|
| Intercept | 63.9 | 0.806 | 79.3 | 527 | < 0.001 |
| Intervention | 0.610 | 1.09 | 0.559 | 527 | 0.576 |
| Female | -2.17 | 1.11 | -1.96 | 527 | 0.051 |
| First-generation | -2.03 | 1.46 | -1.39 | 527 | 0.166 |
| URM | -1.17 | 1.87 | -0.626 | 527 | 0.532 |
| Asian | 0.635 | 1.30 | 0.489 | 527 | 0.625 |
| ACT score | 1.22 | 0.171 | 7.18 | 527 | < 0.001 |
| High school GPA | 9.35 | 1.84 | 5.08 | 527 | < 0.001 |

#### 2.3. Interaction effect between the intervention and URM status

| | Coefficient | Standard error | T-ratio | d.f. | P-value |
|---|---|---|---|---|---|
| Intercept | 64.0 | 0.804 | 79.6 | 526 | < 0.001 |
| Intervention | 0.592 | 1.09 | 0.544 | 526 | 0.587 |
| Female | -2.07 | 1.10 | -1.88 | 526 | 0.061 |
| First-generation | -1.86 | 1.46 | -1.27 | 526 | 0.204 |
| URM | -5.05 | 2.58 | -1.96 | 526 | 0.051 |
| URM*Intervention | 7.73 | 3.55 | 2.18 | 526 | 0.030 |
| Asian | 0.612 | 1.29 | 0.474 | 526 | 0.636 |
| ACT score | 1.23 | 0.170 | 7.21 | 526 | < 0.001 |
| High school GPA | 9.49 | 1.84 | 5.17 | 526 | < 0.001 |

#### Section 3: Study 2 Results

##### 3.1. Performance gaps analysis

Note: control sections only

| | | Coefficient | Error | T-ratio | d.f. | P-value |
|---|---|---|---|---|---|---|
| Level 3 | Intercept | 79.33 | 1.30 | 61.06 | 12 | <.001 |
| Level 1 | Lecture code 1 | -0.90 | 2.61 | -0.35 | 251 | 0.729 |
| | Lecture code 2 | 0.24 | 1.49 | 0.16 | 251 | 0.875 |
| | URM | -3.01 | 1.78 | -1.69 | 251 | 0.092 |
| | MALE | -2.49 | 1.46 | -1.70 | 251 | 0.089 |
| | ACT score | 0.87 | 0.21 | 4.16 | 251 | <.001 |
| | First-generation | -4.13 | 1.75 | -2.36 | 251 | 0.019 |
| | High school GPA | 7.05 | 2.27 | 3.10 | 251 | 0.003 |
| | Asian | -2.27 | 1.88 | -1.21 | 251 | 0.229 |

**LEVEL 1 MODEL** (bold: group-mean centering; bold italic: grand-mean centering)

$$TPERCENT = \pi_0 + \pi_1(LECTX1) + \pi_2(LECTX2) + \pi_3(\boldsymbol{URM}) + \pi_4(MALE) + \pi_5(\boldsymbol{ACT}) + \pi_6(FGEN) + \pi_7(\boldsymbol{HSGPA}) + \pi_8(ASIAN) + e$$

**LEVEL 2 MODEL** (bold: group-mean centering; bold italic: grand-mean centering)

$\pi_0 = \beta_{00} + r_0$

$\pi_1 = \beta_{10} + r_1$

$\pi_2 = \beta_{20} + r_2$

$\pi_3 = \beta_{30} + r_3$

$\pi_4 = \beta_{40} + r_4$

$\pi_5 = \beta_{50} + r_5$

$\pi_6 = \beta_{60} + r_6$

$\pi_7 = \beta_{70} + r_7$

$\pi_8 = \beta_{80} + r_8$

**LEVEL 3 MODEL** (bold italic: grand-mean centering)

$\beta_{0c} = \gamma_{00c} + u_{00}$

$\beta_{1c} = \gamma_{10c} + u_{10}$

$\beta_{2c} = \gamma_{20c} + u_{20}$

$\beta_{3c} = \gamma_{30c} + u_{30}$

$\beta_{4c} = \gamma_{40c} + u_{40}$

$\beta_{5c} = \gamma_{50c} + u_{50}$

$\beta_{6c} = \gamma_{60c} + u_{60}$

$\beta_{7c} = \gamma_{70c} + u_{70}$

$\beta_{8c} = \gamma_{80c} + u_{80}$

#### 3.2 Main effect of intervention and TA mindset

| | | **Coefficient** | **Error** | **T-ratio** | **d.f.** | **P-value** |
|---|---|---|---|---|---|---|
| *Level 3* | Intercept | 79.37 | 1.02 | 77.60 | 14 | <.001 |
| | TA Mindset | 0.70 | 0.42 | 1.64 | 14 | 0.122 |
| *Level 2* | Condition (0=Control; 1=Intervention) | 0.13 | 0.93 | 0.14 | 28 | 0.890 |
| *Level 1* | Lecture code 1 | 0.61 | 1.02 | 0.60 | 565 | 0.549 |
| | Lecture code 2 | 0.06 | 1.10 | 0.05 | 565 | 0.959 |
| | URM | -3.68 | 1.29 | -2.85 | 565 | 0.005 |
| | MALE | -0.86 | 1.21 | -0.71 | 565 | 0.476 |
| | ACT | 0.90 | 0.10 | 8.79 | 565 | <.001 |
| | FGEN | -4.36 | 1.08 | -4.03 | 565 | <.001 |
| | HSGPA | 6.63 | 1.02 | 6.48 | 565 | <.001 |
| | ASIAN | -2.56 | 1.27 | -2.02 | 565 | 0.043 |

**LEVEL 1 MODEL** (bold: group-mean centering; bold italic: grand-mean centering)

TPERCENT = $\pi_0 + \pi_1$(LECTX1) + $\pi_2$(LECTX2) + $\pi_3$(URM) + $\pi_4$(MALE) + $\pi_5$(***ACT***) + $\pi_6$(FGEN) + $\pi_7$(***HSGPA***) + $\pi_8$(ASIAN) + $e$

**LEVEL 2 MODEL** (bold: group-mean centering; bold italic: grand-mean centering)

$\pi_0 = \beta_{00} + \beta_{01}$(CONDITIO) + $r_0$

$\pi_1 = \beta_{10} + r_1$

$\pi_2 = \beta_{20} + r_2$

$\pi_3 = \beta_{30} + r_3$

$\pi_4 = \beta_{40} + r_4$

$\pi_5 = \beta_{50} + r_5$

$\pi_6 = \beta_{60} + r_6$

$\pi_7 = \beta_{70} + r_7$

$\pi_8 = \beta_{80} + r_8$

**LEVEL 3 MODEL** (bold italic: grand-mean centering)

$\beta_{00} = \gamma_{000} + \gamma_{001}$(***MINDSET***) + $u_{00}$

$\beta_{01} = \gamma_{010} + u_{01}$

$\beta_{10} = \gamma_{100} + u_{10}$

$\beta_{20} = \gamma_{200} + u_{20}$

$\beta_{30} = \gamma_{300} + u_{30}$

$\beta_{40} = \gamma_{400} + u_{40}$

$\beta_{50} = \gamma_{500} + u_{50}$

$\beta_{60} = \gamma_{600} + u_{60}$

$\beta_{70} = \gamma_{700} + u_{70}$

$\beta_{80} = \gamma_{800} + u_{80}$

##### 3.3 Interaction effect between the intervention and first generation status

| | Effect | Coefficient | Error | T-ratio | d.f. | P-value |
|---|---|---|---|---|---|---|
| *Level 3* | Intercept | 79.18 | 1.05 | 75.56 | 14 | <.001 |
| | TA Mindset | 0.69 | 0.43 | 1.61 | 14 | 0.130 |
| *Level 2* | Condition (0=Control; 1=Intervention) | 0.46 | 0.93 | 0.50 | 28 | 0.620 |
| *Level 1* | Lecture code 1 | 0.58 | 1.03 | 0.56 | 564 | 0.576 |
| | Lecture code 2 | 0.06 | 1.09 | 0.06 | 564 | 0.957 |
| | URM | -3.69 | 1.30 | -2.83 | 564 | 0.005 |
| | MALE | -0.86 | 1.22 | -0.71 | 564 | 0.481 |
| | ACT | 0.91 | 0.10 | 8.84 | 564 | <.001 |
| | First-generation | -3.62 | 0.99 | -3.64 | 564 | 0.001 |
| | First generation x Condition | -1.35 | 1.21 | -1.11 | 564 | 0.266 |
| | HSGPA | 6.64 | 1.02 | 6.54 | 564 | <.001 |
| | ASIAN | -2.52 | 1.28 | -1.97 | 564 | 0.048 |

**LEVEL 1 MODEL** (bold: group-mean centering; bold italic: grand-mean centering)

$$TPERCENT = \pi_0 + \pi_1(LECTX1) + \pi_2(LECTX2) + \pi_3(URM) + \pi_4(MALE) + \pi_5(\boldsymbol{ACT}) + \pi_6(FGEN) + \pi_7(\boldsymbol{HSGPA}) + \pi_8(ASIAN) + e$$

**LEVEL 2 MODEL** (bold: group-mean centering; bold italic: grand-mean centering)

$$\pi_0 = \beta_{00} + \beta_{01}(CONDITIO) + r_0$$

$$\pi_1 = \beta_{10} + r_1$$

$$\pi_2 = \beta_{20} + r_2$$

$$\pi_3 = \beta_{30} + r_3$$

$$\pi_4 = \beta_{40} + r_4$$

$$\pi_5 = \beta_{50} + r_5$$

$$\pi_6 = \beta_{60} + \beta_{61}(CONDITIO) + r_6$$

$$\pi_7 = \beta_{70} + r_7$$

$$\pi_8 = \beta_{80} + r_8$$

**LEVEL 3 MODEL** (bold italic: grand-mean centering)

$$\beta_{00} = \gamma_{000} + \gamma_{001}(\boldsymbol{MINDSET}) + u_{00}$$

$$\beta_{01} = \gamma_{010} + u_{01}$$

$$\beta_{10} = \gamma_{100} + u_{10}$$

$$\beta_{20} = \gamma_{200} + u_{20}$$

$$\beta_{30} = \gamma_{300} + u_{30}$$

$$\beta_{40} = \gamma_{400} + u_{40}$$

$$\beta_{50} = \gamma_{500} + u_{50}$$

$$\beta_{60} = \gamma_{600} + u_{60}$$

$$\beta_{61} = \gamma_{610} + u_{61}$$

$$\beta_{70} = \gamma_{700} + u_{70}$$

$$\beta_{80} = \gamma_{800} + u_{80}$$

#### 3.4 Interaction effect between the intervention and TA mindset

| | | Coefficient | Error | T-ratio | d.f. | P-value |
|---|---|---|---|---|---|---|
| *Level 3* | Intercept | 79.34 | 1.00 | 79.19 | 14 | <.001 |
| | TA Mindset | 1.47 | 0.49 | 3.02 | 14 | 0.01 |
| *Level 2* | Condition (0=Control; 1=Intervention) | 0.27 | 0.88 | 0.31 | 28 | 0.758 |
| | TA Mindset x Condition | -1.44 | 0.36 | -4.01 | 28 | 0.001 |
| *Level 1* | Lecture code 1 | 0.12 | 0.93 | 0.13 | 564 | 0.896 |
| | Lecture code 2 | 0.40 | 0.97 | 0.41 | 564 | 0.684 |
| | URM | -3.71 | 1.33 | -2.80 | 564 | 0.006 |
| | MALE | -0.80 | 1.18 | -0.68 | 564 | 0.496 |
| | ACT score | 0.90 | 0.10 | 8.99 | 564 | <.001 |
| | First-generation | -4.42 | 1.10 | -4.01 | 564 | <.001 |
| | High school GPA | 6.51 | 1.07 | 6.11 | 564 | <.001 |
| | Asian | -2.59 | 1.26 | -2.06 | 564 | 0.039 |

**LEVEL 1 MODEL** (bold: group-mean centering; bold italic: grand-mean centering)

TPERCENT $= \pi_0 + \pi_1$(LECTX1) $+ \pi_2$(LECTX2) $+ \pi_3$(URM) $+ \pi_4$(MALE) $+ \pi_5$(***ACT***) $+ \pi_6$(FGEN) $+ \pi_7$(***HSGPA***) $+ \pi_8$(ASIAN) $+ e$

**LEVEL 2 MODEL** (bold: group-mean centering; bold italic: grand-mean centering)

$\pi_0 = \beta_{00} + \beta_{01}$(CONDITIO) $+ r_0$

$\pi_1 = \beta_{10} + r_1$

$\pi_2 = \beta_{20} + r_2$

$\pi_3 = \beta_{30} + r_3$

$\pi_4 = \beta_{40} + r_4$

$\pi_5 = \beta_{50} + r_5$

$\pi_6 = \beta_{60} + r_6$

$\pi_7 = \beta_{70} + r_7$

$\pi_8 = \beta_{80} + r_8$

**LEVEL 3 MODEL** (bold italic: grand-mean centering)

$\beta_{00} = \gamma_{000} + \gamma_{001}$(***MINDSET***) $+ u_{00}$

$\beta_{01} = \gamma_{010} + \gamma_{011}$(***MINDSET***) $+ u_{01}$

$\beta_{10} = \gamma_{100} + u_{10}$

$\beta_{20} = \gamma_{200} + u_{20}$

$\beta_{30} = \gamma_{300} + u_{30}$

$\beta_{40} = \gamma_{400} + u_{40}$

$\beta_{50} = \gamma_{500} + u_{50}$

$\beta_{60} = \gamma_{600} + u_{60}$

$\beta_{70} = \gamma_{700} + u_{70}$

$\beta_{80} = \gamma_{800} + u_{80}$
